# Mevalonate pathway-mediated ER homeostasis is required for haploid stability in human somatic cells

**DOI:** 10.1101/2020.11.05.369231

**Authors:** Kan Yaguchi, Kimino Sato, Koya Yoshizawa, Gabor Banhegyi, Eva Margittai, Ryota Uehara

## Abstract

The somatic haploidy is unstable in diplontic animals, but cellular processes determining haploid stability remain elusive. Here, we found that inhibition of mevalonate pathway by pitavastatin, a widely used cholesterol-lowering drug, drastically destabilized the haploid state in HAP1 cells. Interestingly, cholesterol supplementation did not restore haploid stability in pitavastatin-treated cells, and cholesterol inhibitor U18666A did not phenocopy haploid destabilization. These results ruled out the involvement of cholesterol in haploid stability. Besides cholesterol perturbation, pitavastatin induced endoplasmic reticulum (ER) stress, the suppression of which by a chemical chaperon significantly restored haploid stability in pitavastatin-treated cells. Our data demonstrate the involvement of the mevalonate pathway in the stability of the haploid state in human somatic cells through managing ER stress, highlighting a novel link between ploidy and ER homeostatic control.

## Introduction

Ploidy alteration is a hallmark of cancers. While tetraploidy occurring through whole-genome duplication is the most prevalent type of ploidy alteration in solid tumors (Bielski *et al*., 2018; Carter *et al.*, 2012; Dewhurst *et al.*, 2014), near-haploidy is also seen in certain rare types of blood and solid cancers with poor prognosis (Brodeur *et al*., 1981; Kohno *et al*., 1980; Oshimura *et al*., 1977; Safavi *et al*., 2013; Safavi and Paulsson, 2017; Sukov *et al*., 2010). In diplontic animal organisms, somatic haploidy is generally unstable, causing frequent autodiploidization at the cellular level and severe developmental abnormalities at the organismal level (Sagi and Benvenisty, 2017; Wutz, 2014).

The halving of genome copy from the normal diploid state potentially has pleiotropic effects on cellular homeostasis in haploid cells. An apparent feature of haploid cells is their halved cellular volume to diploids with the halving of total protein content (Yaguchi *et al.*, 2018). Though these features possibly have profound influence on intracellular processes – such as the metabolic control – in haploid state, it remains elusive what aspects of the metabolism alter and characterize cellular phenotypes of haploid cells.

The mevalonate pathway metabolizes acetyl-CoA to produce sterol isoprenoids, and non-sterol isoprenoids that mediate diverse biosynthetic processes essential for cell construction and proliferation (Buhaescu and Izzedine, 2007; Mullen *et al*., 2016). Among mevalonate-derived metabolites, cholesterol serves as a structural component of cell membranes and a precursor of fundamental biomolecules, such as steroid hormones. Mevalonate-derived polyisoprenols, such as dolichol phosphates are essential components of glycoprotein synthesis and endoplasmic reticulum (ER) homeostasis participating in protein N-glycosylation, C- and O-mannosylation, and GPI-anchor production (Carlberg *et al*., 1996; Chojnacki and Dallner, 1988; Doucey *et al*., 1998). Mevalonate-derived isoprenoids are also used for the prenylation of small GTPases, which mediates their association to the cell membrane and signal transduction for dynamic processes such as cytoskeletal reorganization and vesicular trafficking (Leung *et al.*, 2006; Wang and Casey, 2016). Recent studies have revealed that the mevalonate pathway controls cell size by optimizing mitochondrial functionality through protein prenylation (Miettinen and Björklund, 2015; Miettinen and Björklund, 2016; Miettinen *et al*., 2014). Inhibition of the rate-limiting enzyme of mevalonate pathway, 3-hydroxy-3-methylglutaryl-coenzyme A reductase (HMGCR) by statins perturbs this homeostatic control, leading to increased cell size in different types of cultured cells (Miettinen and Björklund, 2015).

In this study, in an attempt to modulate cell size by an HMGCR inhibitor pitavastatin in human haploid HAP1 cells (Carette *et al*., 2011), we found that the inhibitor compromised the stability of the haploid state in these cells. Interestingly, a recent chemical screen searching for compounds that stabilize haploid state has also identified statins leading to the selective loss of haploid cells (Olbrich *et al*., 2019). However, whether the mevalonate pathway is indeed involved in promoting haploid stability, and how the inhibition of the pathway may lead to destabilization of haploid state is still unknown. Using a pharmacological approach, we specified that statin-induced ER stress as a process responsible for the destabilization of the haploid state.

## Materials and methods

### Cell culture and flow cytometry

Near haploid human cell line, HAP1 (Carette et al., 2011) was purchased from Haplogen and cultured in Iscove’s Modified Dulbecco’s Medium (IMDM; Wako Pure Chemical Industries) supplemented with 10% fetal bovine serum and 1× antibiotic-antimycotic (Sigma-Aldrich). Haploid cells were regularly maintained by size-based cell sorting, as previously described (Yaguchi et al., 2018). For DNA content analysis, cells were stained with 10 μg/ml Hoechst 33342 (Dojindo) for 15 min at 37°C, and fluorescence intensity was analyzed using a JSAN desktop cell sorter (Bay bioscience).

### Chemical compounds

Compounds were purchased from suppliers as follows: Pitavastatin (163-24861, Wako); U18666A (10009085, Cayman Chemical); Cholesterol (SLBZ0657, Sigma-Aldrich); FTI-277 (S7465, Selleck); GGTI-298 (S7466, Selleck); tauroursodeoxycholic acid (TUDCA, T1567, Tokyo Chemical Industry); and mevalonate (mevalonolactone, M4667, Sigma-Aldrich). Cholesterol was visualized using the Cholesterol Cell-Based Detection Assay Kit (10009779, Cayman Chemical) according to the manufacture’s instruction.

### Long-term cell passage experiments

Freshly purified haploid HAP1 cells were cultured in the presence of different compounds at final concentrations described elsewhere. We conducted cell passage using 0.05% trypsin-EDTA (Wako), typically once two days. To investigate the effect of each compound on the stability of haploid state, we continued passages for ~3 weeks before subjecting cell culture to flow cytometric DNA content analysis. As an exception, we conducted DNA content analysis after ~1-week passages for FTI-277-treated culture because drastic cell death precluded further passages. As an index of haploid cell preservation, we quantified the proportion of haploid cells in the G1 cell cycle phase (haploid G1 proportion) in flow cytometric analysis.

### Antibodies and immunoblotting

Antibodies were purchased from suppliers and used at dilutions, as follows: mouse monoclonal anti-CHOP (L63F7, Cell Signaling technology; 1:1000); rabbit monoclonal anti-ATF4 (D48B, Cell signaling technology; 1:1000); mouse monoclonal anti-β-tubulin (10G10, Wako; 1:1000). Horseradish peroxidase-conjugated secondary antibodies were purchased from Jackson ImmunoResearch Laboratories and used at a dilution of 1:1000. Immunoblotting was performed as previously described (Yaguchi et al., 2018). We used the ezWestLumi plus ECL Substrate (ATTO) and a LuminoGraph II chemiluminescent imaging system (ATTO) for signal detection. We quantified immunoblotting signals using the Gels tool in ImageJ software (National Institutes of Health).

## Results

### Mevalonate production is required for the stability of the haploid state in HAP1 cells

To test the effects of inhibition of the mevalonate pathway on cell size control, we treated human haploid cell HAP1 with a competitive HMGCR inhibitor pitavastatin, which has been reported to increase the size of human cell lines such as Jurkat (Miettinen and Björklund, 2015). Although we did not observe the increase of cell size in HAP1 cells upon pitavastatin treatment (Fig. 1A), we unexpectedly found that the inhibitor destabilized the haploid state of HAP1 cells in flow cytometric DNA content analysis (Fig. 1B and C). Freshly purified, non-treated haploid HAP1 cells gradually diploidized during ~3-weeks continuous passages, resulting in the reduction in haploid G1 proportion to 28.0 ± 1.7% (mean ± standard error, n =3; Fig. 1B and C) (Yaguchi et al., 2018). However, when treated with 0.5 μM pitavastatin, which allowed long-term culture without blocking cell proliferation, the progression of haploid-to-diploid conversion was substantially accelerated (haploid G1 proportion of 17.1 ± 1.2%; Fig. 1B and C). Since this result indicates the importance of statin-targeted processes in haploid stability, we further investigated the mechanism underlying the statin-mediated destabilization of haploid state. First, we tested whether co-treatment with mevalonate ameliorates haploid stability in the presence of pitavastatin (Fig. 1B). Mevalonate supplementation significantly preserved the haploid population in pitavastatin-treated culture (haploid G1 proportion of 30.2 ± 1.0%; Fig. 1C), demonstrating the requirement for sufficient amount of mevalonate for haploid stability in HAP1 cells.

**Fig. 1:**
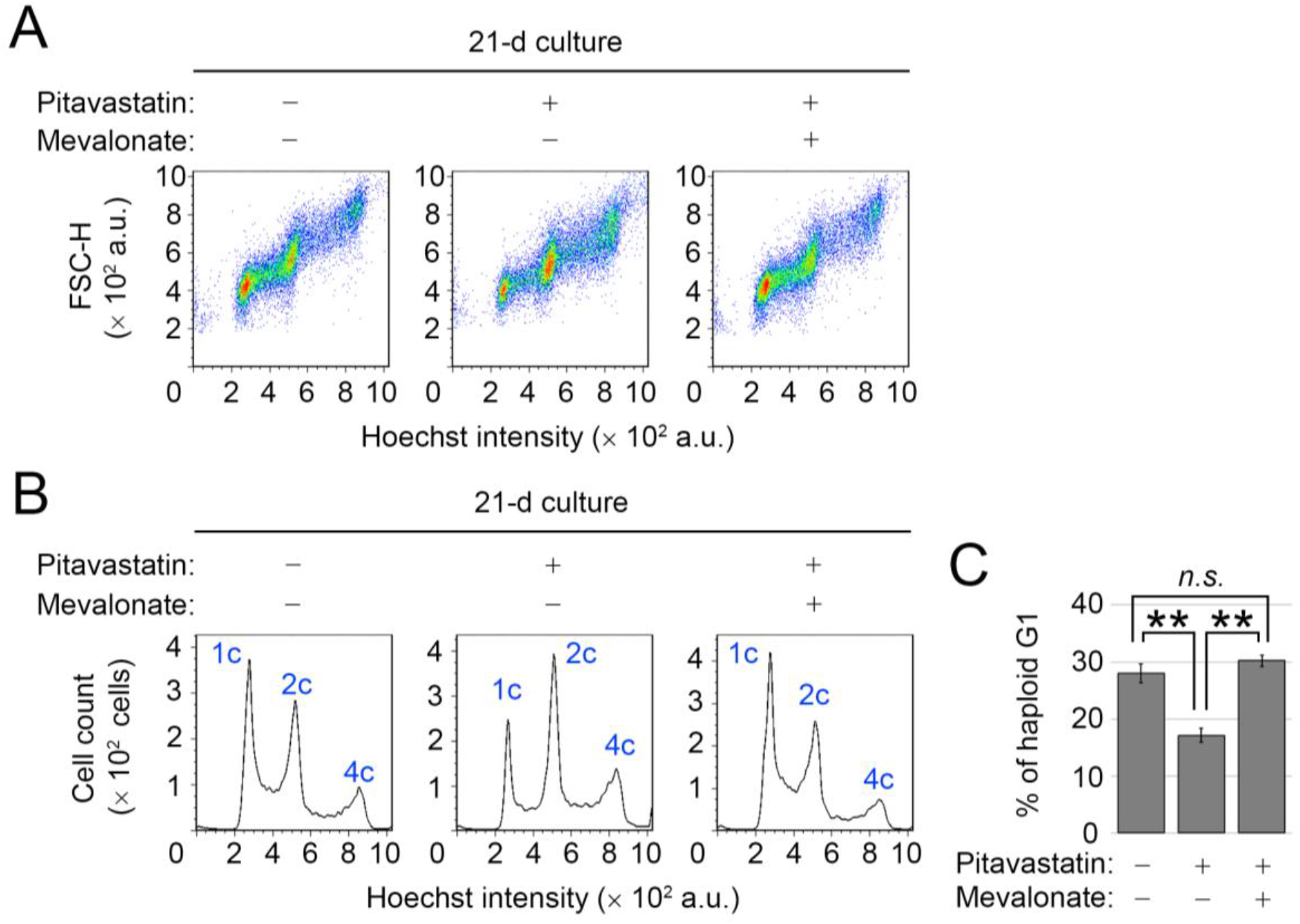
Destabilization of the haploid state by inhibition of the mevalonate pathway in HAP1 cells. (**A, B**) Flow cytometric analysis of DNA content in Hoechst-stained cells after 21-d culture. Cells were cultured in the absence or presence of 0.5 μM pitavastatin with or without 20 μM mevalonate supplementation. Dot plots of forward scatter signal (for the judgment of relative cell size) against Hoechst signal (A) and histograms of Hoechst signal (B). (**C**) The proportion of the haploid G1 population in B. Means ± standard error (SE) of three independent experiments (day 20 or 21 in the long-term passages, \*\**p*< 0.01, n.s.: not significant, oneway ANOVA with Tukey post-hoc test).

### Cholesterol perturbation is not the cause of haploid destabilization by pitavastatin

Next, we determined downstream branches of the mevalonate pathway crucial for the maintenance of haploid state. Because statins are widely used cholesterol-lowering drugs (Adhyaru and Jacobson, 2018), we addressed the possible involvement of the cholesterol branch in haploid stability. For this, we visualized intracellular distribution and content of cholesterol in control and pitavastatin-treated HAP1 cells using a cholesterol-binding fluorescent compound filipin (Fig. 2A) (Drabikowski *et al.*, 1973). In control cells, filipin fluorescence signal distributed throughout the plasma- and intracellular membrane structures, and 0.5 μM pitavastatin modestly, but significantly, reduced the filipin staining intensity (Fig. 2B and C). Therefore, at this final concentration, pitavastatin mildly reduced cholesterol level in HAP1 cells. Next, we addressed whether cholesterol supplementation is sufficient to restore haploid stability in pitavastatin-treated cells. The addition of 10 μM cholesterol to the pitavastatin-treated cell culture fully restored cholesterol level visualized by filipin staining (Fig. 2A-C). However, cholesterol supplementation did not affect the progression of haploid-to-diploid conversion in statin-treated culture (Fig. 2D and E). On the other hand, mevalonate supplementation, which fully restored haploid stability in statin-treated cells (Fig. 1), did not change the cholesterol level in statin-treated cells (Fig. 2C). These data demonstrated that lowered cholesterol level was not the cause of haploid destabilization by pitavastatin.

**Fig. 2:**
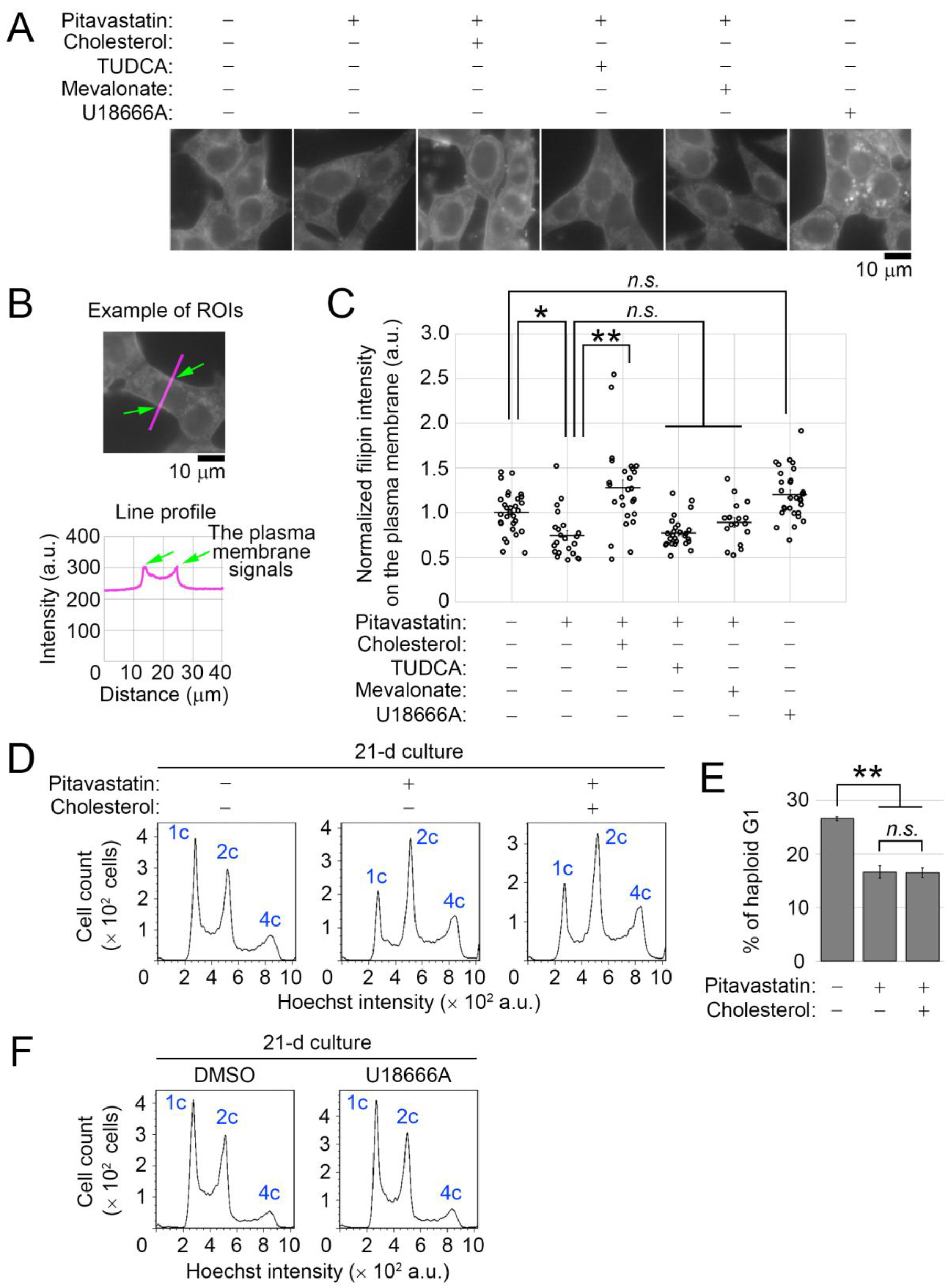
Either supplementation or perturbation of cholesterol does not affect haploid stability. (**A**) Fluorescence microscopy of HAP1 cells stained by filipin after treating with the compounds for 1 d. (**B, C**) Quantification of filipin fluorescence intensity on the plasma membrane in A. The fluorescence signals on the plasma membrane were quantified from line profiles taken across the cells, as shown in B. Means ± SE of at least 18 cells from two independent experiments (\**p* < 0.05, \*\**p* < 0.01, one-way ANOVA with Tukey post-hoc test). (**D**) DNA content analysis after 21-d culture. Cells were cultured in the absence or presence of 0.5 μM pitavastatin with or without 10 μM cholesterol supplementation. (**E**) The proportion of the haploid G1 population in D. Means ± SE of three independent experiments (day 21 in the long-term passages, \*\**p*< 0.01, one-way ANOVA with Tukey post-hoc test). (**F**) DNA content analysis after 21-d culture. Cells were cultured in the absence or presence of 2.5 μM U18666A. Representative data from two independent experiments.

Next, we tested the effect of perturbation of cholesterol homeostasis by a non-statin cholesterol inhibitor on haploid stability. An amphipathic steroid U18666A perturbs the cholesterol-mediated bioprocesses by inhibiting both synthesis and intracellular transport of cholesterol (Cenedella, 2009). Treatment with 2.5 μM U18666A resulted in the accumulation of cholesterol in intracellular vesicles, a typical defect caused by the compound (Fig. 2A) (Reiners *et al*., 2011; Underwood *et al*., 1998). HAP1 cell proliferation was not severely affected by 2.5 μM U18666A, allowing us to test its effect on the long-term haploid stability. In the long-term passages, 2.5 μM U18666A-treated cells underwent haploid-to-diploid conversion at a similar pace as non-treated control (Fig. 2F). This result further ruled out the possible involvement of cholesterol homeostatic control in haploid stability in HAP1 cells.

### Inhibition of protein prenylation does not phenocopy haploid destabilization by pitavastatin

Among the mevalonate-derived metabolites, farnesyl pyrophosphate and geranylgeranyl pyrophosphate are used for the posttranslational prenylation of small GTPases that play crucial roles in the regulation of cell cycle and proliferation, as well as cell size control (Berndt *et al.*, 2011; Miettinen and Björklund, 2015; Miettinen and Björklund, 2016). Therefore, we tested the effects of FTI-277 or GGTI-298, which inhibits protein farnesylation or geranylgeranylation, respectively, on the stability of the haploid state in HAP1 cells. Twenty μM FTI-277 treatment caused mitotic progression defects marked by the round-shaped mitotically-arrested cells and abnormally enlarged cells in culture (Fig. 3A). Similar FTI-277-induced mitotic defects have been reported in different cell lines (Holland *et al*., 2015; Morgan *et al*., 2001; Moudgil *et al*., 2015). Consistent with the microscopic observation, FTI-277-treated HAP1 cells were drastically polyploidized within several days with the prominent accumulation of 2, 4, and 8 c peaks in flow cytometric analysis (Fig. 3B). This result suggests that FTI-277 induces whole-genome duplication in HAP1 cells regardless of the ploidy state, which was in contrast to the haploidy-specific induction of whole-genome duplication by pitavastatin. The drastic polyploidization and subsequent cell death precluded us from testing the effects of FTI-277 on ploidy dynamics in a more extended period.

**Fig. 3:**
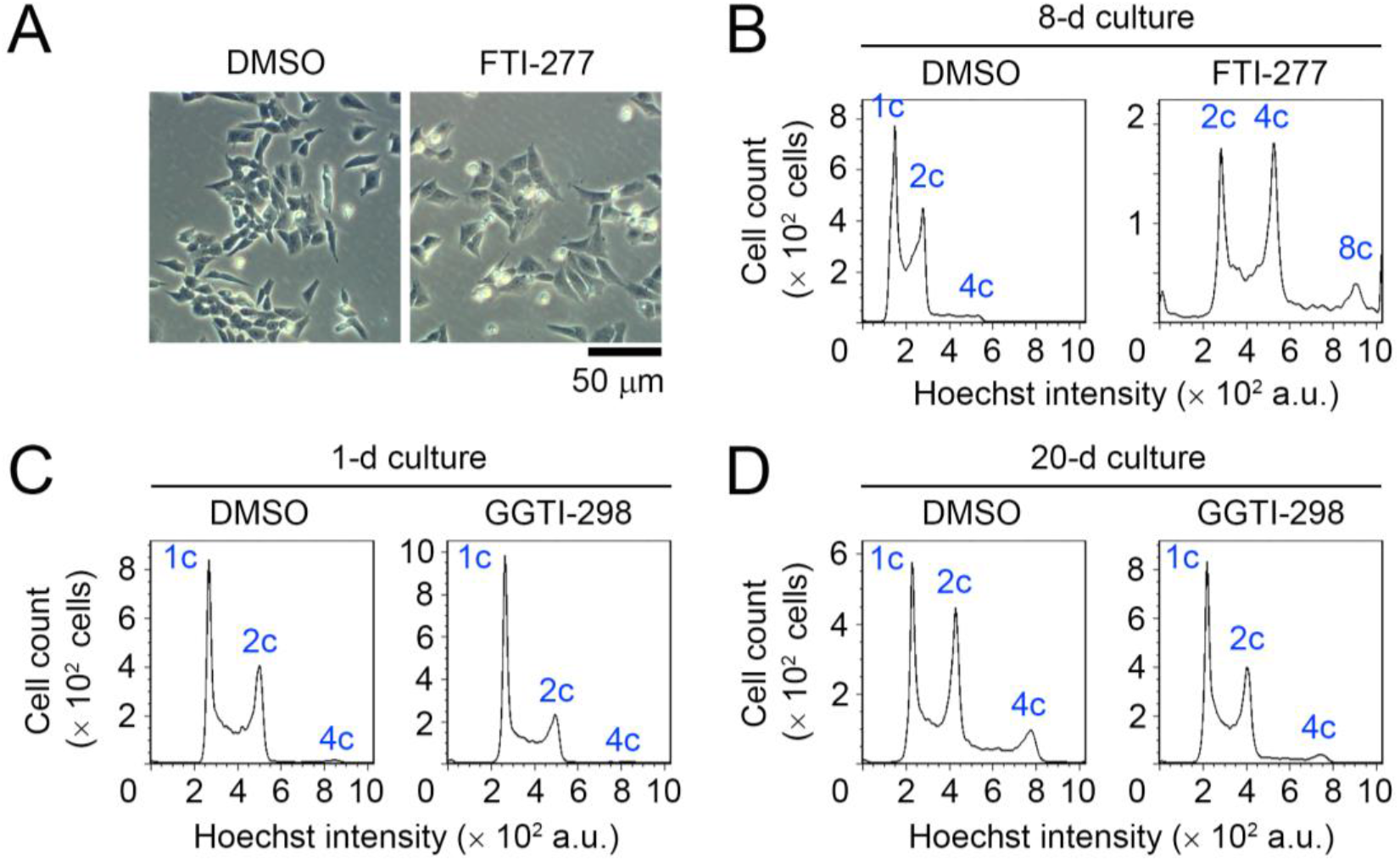
Inhibition of protein prenylation does not phenocopy the statin-induced haploid destabilization. (**A**) Transparent microscopy of HAP1 cells treated with or without 20 μM FTI-277 for 2 d. Representative data from two independent experiments. (**B-D**) DNA content analysis after 8-d (B), 1-d (C), or 20-d culture (D). Cells were cultured with or without 20 μM FTI-277 (B) or 2 μM GGTI-298 (C, D). Representative data from two independent experiments.

On the other hand, treatment with 2 μM GGTI-298 mildly arrested haploid HAP1 cells at the G1 phase within 24 h (Fig. 3C), consistent with a previous report in several cell types (Sun *et al*., 1999). In prolonged culture for 20 d in the presence of 2 μM GGTI-298, the haploid-to-diploid conversion was considerably slowed down compared to non-treated control, presumably because of the moderate G1 arrest (Fig. 3D). Therefore, the suppression of either protein farnesylation or geranylgeranylation did not phenocopy the pitavastatin-induced haploid destabilization in our long-term experiment.

### Pitavastatin destabilizes the haploid state by evoking ER stress

Since statins potentially induce ER stress by suppressing dolichol phosphates biosynthesis and inhibiting protein N-glycosylation (Chojnacki and Dallner, 1988), we next tested the possibility that pitavastatin destabilizes the haploid state through perturbing ER homeostasis. For this, we tested the effect of pitavastatin on ER stress in HAP1 cells using immunoblot analysis of ATF4 or CHOP/GADD153/DDIT3, the unfolded protein response (UPR) components whose expression increases upon the induction of ER stress (Iurlaro and Muñoz-Pinedo, 2016; Marciniak *et al.*, 2004; Wang *et al*., 2019). Treatment with 0.5 μM pitavastatin for 3 d significantly increased the expression of both ATF4 and CHOP (Fig. 4A and B). Mevalonate supplementation canceled the ATF4 and CHOP upregulation in pitavastatin-treated cells (Fig. 4A and B), demonstrating that pitavastatin evoked ER stress specifically through blocking mevalonate metabolism.

**Fig. 4:**
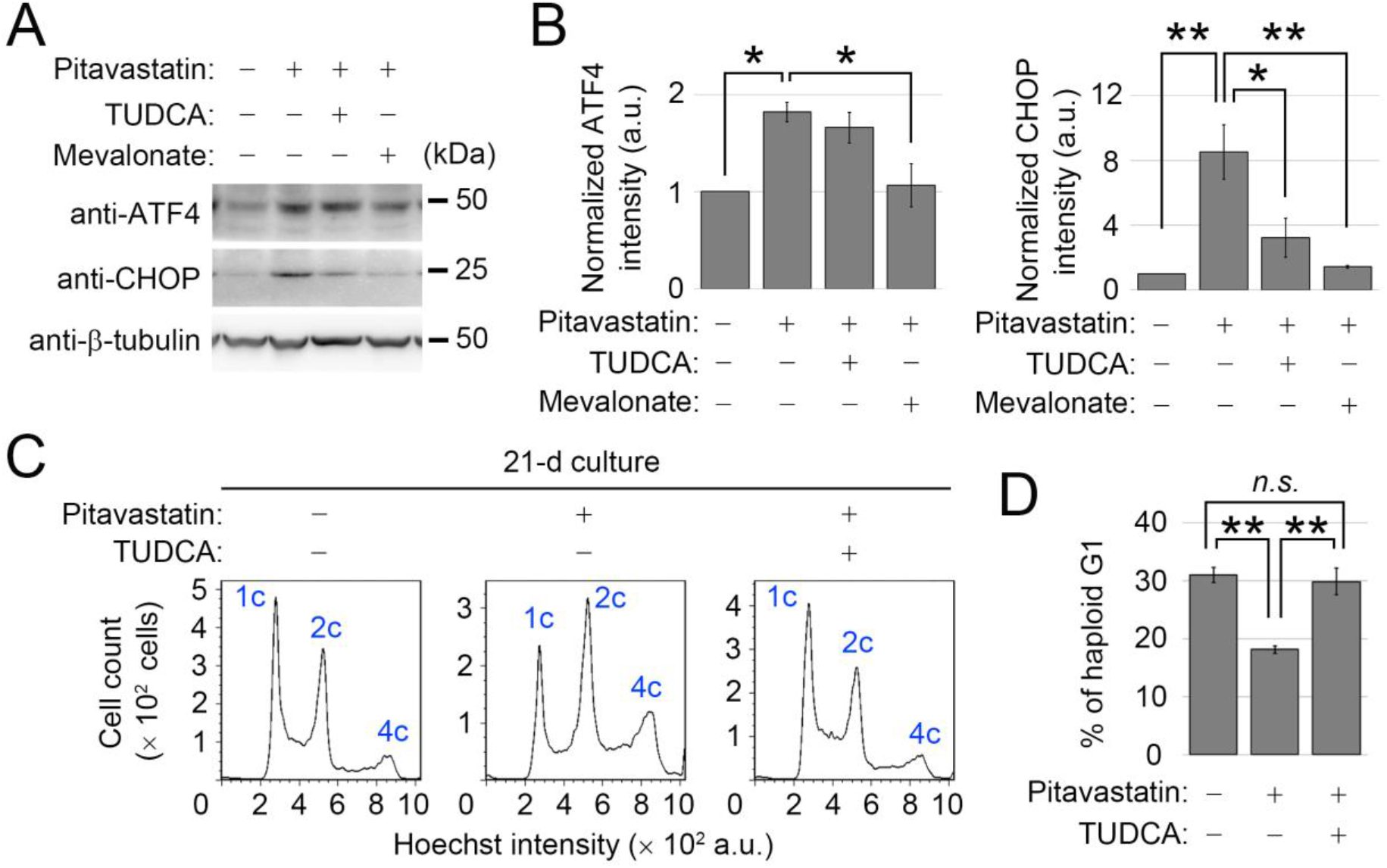
Amelioration of ER stress improves haploid stability in pitavastatin-treated cells. (**A**) Immunoblotting of ATF4 and CHOP in HAP1 cells treated with the compounds for 3 d. β-tubulin was detected as a loading control. (**B**) Quantification of relative expression of ATF4 and CHOP in A. Means ± SE of three independent experiments (\**p* < 0.05, \*\**p* < 0.01, one-way ANOVA with Tukey post-hoc test). (**C**) DNA content analysis after 21-d culture. Cells were cultured in the absence or presence of 0.5 μM pitavastatin with or without 2.5 mM TUDCA. (**D**) The proportion of the haploid G1 population in C. Means ± SE of three independent experiments (day 21 in the long-term passages, \*\**p* < 0.01, one-way ANOVA with Tukey post-hoc test).

Finally, we determined whether ER stress induction is the cause of pitavastatin-mediated destabilization of haploid state in HAP1 cells. For this, we tested the effect of an ER stress-reducing chemical chaperone, tauroursodeoxycholic acid (TUDCA) (Ozcan *et al.*, 2006; Yoon *et al*., 2016), on the haploid stability of HAP1 cells. Co-treatment with TUDCA did not affect ATF4 expression, but substantially blocked CHOP upregulation in pitavastatin-treated cells (Fig. 4A and B), presumably reflecting the complex effects of chemical chaperones on different factors in the UPR pathways (Uppala *et al*., 2017). In contrast, TUDCA did not change the cholesterol level in pitavastatin-treated cells assessed by filipin staining (Fig. 2A-C). In long-term passages, co-treatment of TUDCA significantly slowed down haploid-to-diploid conversion in pitavastatin-treated cells (Fig. 4C and D). Therefore, restoration of ER homeostasis by TUDCA substantially improved the stability of the haploid state in the presence of pitavastatin, demonstrating that haploid destabilization by pitavastatin is caused, at least in part, through the induction of ER stress.

## Discussion

It is assumed that ploidy differences have pleiotropic effects on intracellular biosynthetic processes and that the altered biosynthesis, in turn, affects cellular physiology at different ploidy states. However, it remains mostly elusive what biosynthetic processes have influences on ploidy-linked cellular phenotypes. In this study, we found that an HMGCR inhibitor pitavastatin destabilized haploid state in human HAP1 cells. This result is consistent with the recent compound screen that identified statins to efficiently promote the expansion of diploidized population over haploids in HAP1 cell culture (Olbrich et al., 2019). As statins are widely used cholesterol-lowering drugs, the possible involvement of cholesterol metabolism in haploid stability has been suggested in the previous study (Olbrich et al., 2019). Interestingly, however, our results in the current study exclude this possibility for three reasons; 1) full restoration of cholesterol level by cholesterol supplementation did not improve haploid stability in statin-treated cells, 2) mevalonate supplementation fully restored haploid stability without restoring cholesterol level in statin-treated cells, and 3) cholesterol perturbation by a non-statin compound did not affect haploid stability.

Our data further specified the perturbation of mevalonate-mediated ER homeostatic control as a critical cause of the statin-induced haploid destabilization. Interestingly, pitavastatin-induced ER stress caused a drastic acceleration of haploid-to-diploid conversion without affecting the stability of diploid state in HAP1 cells. It remains unknown why the effect of pitavastatin on genome stability was specific to the haploid state. However, a possible reason might be a ploidy-dependent difference in tolerance to ER stress. Haploid cells are half in cell volume than diploid cells (Yaguchi et al., 2018), which presumably restricts intracellular spatial capacity for organelle structures. It has been demonstrated that the expansion of the ER lumen serves as a mechanism to increase ER capacity to ameliorate ER stress upon the accumulation of unfolded proteins (Bernales *et al*., 2006; Schuck *et al*., 2009; Shaffer *et al*., 2004; Sriburi *et al*., 2004). The lower availability of intracellular space may limit stress-responding ER expansion in haploid cells, hence lower tolerance to unfolded protein accumulation.

Mevalonate metabolism is an essential process that supports diverse biosynthetic pathways. Though we specified the preservation of ER homeostasis as a critical process underlying statin-mediate haploid destabilization, we cannot rule out other mevalonate-derived biosynthetic processes in haploid stability. For example, we cannot exclude the possibility that specific targets of protein farnesylation play roles in haploid stability, which might have been masked by the gross polyploidization upon the treatment with FTI-277. Comparative metabolome analysis would be a powerful approach to elucidate other biosynthetic processes playing critical roles in determining the physiology of cells at different ploidy states.

## Acknowledgment

This work was supported by Grant-in-Aid for JSPS Fellows to K.Ya (Grant #19J12210), the Hungarian National Research, Development and Innovation Office (NKFIH grant number: FK124442) and János Bolyai Research Scholarship from the Hungarian Academy of Sciences to E.M., Grants-in-Aid for Scientific Research B (19H03219) and on Innovative Areas “Singularity Biology (No.8007)” (19H05413), and Fostering Joint International Research B (19KK0181) of MEXT, the Princess Takamatsu Cancer Research Fund, the Kato Memorial Bioscience Foundation, the Orange Foundation, the Smoking Research Foundation, the Suhara Memorial Foundation, and the Nakatani Foundation to R.U., and Bilateral Joint Research Projects of Japan Society for the Promotion of Science and Hungarian Academy of Sciences (JPJSBP120193801) to G.B., E.M., and R.U.,

## Author Contributions

Conceptualization, K.Ya., and R.U.; Methodology, K.Ya., K. S., G.B., E.M., and R.U.; Investigation, K.Ya, K.S. K.Yo., E.M., and R.U.; Formal Analysis, K.Ya., K.S., and R.U.; Resources, G.B., E.M., and R.U.; Writing – Original Draft, K.Ya., and R.U.; Writing – Review & Editing, K.Ya., E.M., and R.U.; Funding Acquisition, K.Ya., G.B., E.M., and R.U.

